## Supplementary information for "Optically detected and radio wave-controlled spin chemistry in flavoproteins"

#### Supplementary Note 1. MFE Laser power dependence of *Cra*Cry

The dependence of the magnetic field effect on laser power was characterized over a range of 0.12 to 5.92 μW μm$^{-2}$ (10 linear steps). The overall dynamics toward the new photochemical equilibrium after photoexcitation were subtracted from the dataset shown in Figure S1. The MFE contrast remains approximately constant across different laser powers, revealing that photon flux has only minor impact on the *Cra*Cry MFE.

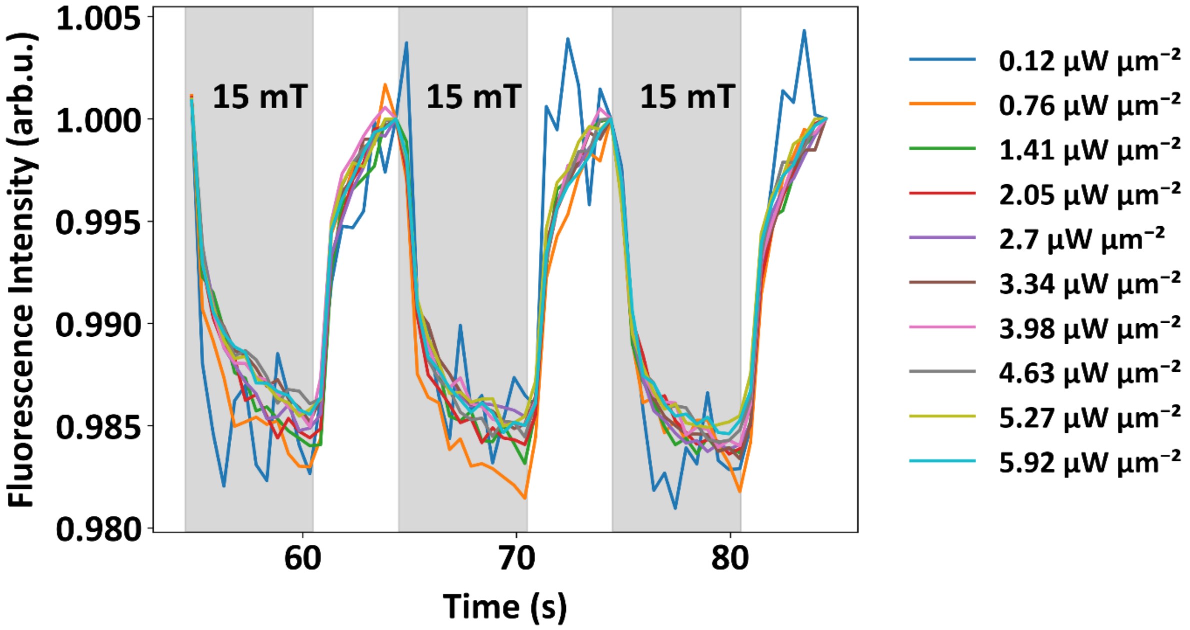


**Figure S1. MFE laser power dependence of *Cra*Cry.**

#### Supplementary Note 2. Reference experiments

We observe a negative magnetic field effect—that is, a reduction in fluorescence intensity upon application of a magnetic field in our *Cra*Cry data. This contrasts with previous studies involving a different class of cryptochromes[1] (e.g., *At*Cry). Although the underlying reasons are not yet understood, we currently attribute this difference to experimental conditions (e.g., optical excitation power and fluorescence filters) and/or differences in decay pathways and kinetics between the cryptochrome classes. Future work will further investigate these findings.

In order to exclude the influence of free FAD in solution, which is known to cause a negative field effect, we performed additional experiments[1,2]. Each experiment was performed with a fresh thawed sample of *Cra*Cry, to avoid degradation. UV/Vis spectra were recorded for checking the protein state, which shows characteristic features of an intact protein (Fig. S2). The most prominent features of bound FAD are a slight red shift of the absorption maximum from 445 to 450 nm and the three-peak pattern of this absorption maximum[3,4], both of which are clearly observed in our samples.

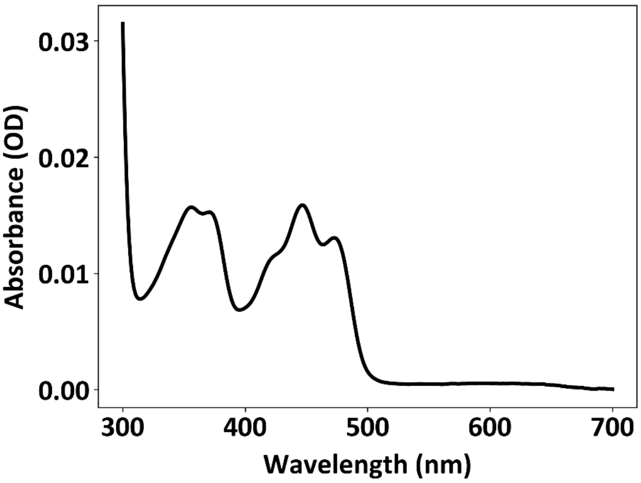


**Figure S2. Typical UV-Vis spectra of a representative *Cra*Cry sample.** Protein sample directly taken from -80 °C and prepared as described in the sample preparation section before ODMR or MFE measurements.

Degradation of the protein and escape of the flavin to the solution is clearly observed after long measurements (which we avoided). We purposely exposed *Cra*Cry to 37° for 16 hours and observed that the ODMR signal disappeared, which proves that an intact protein is needed for efficient ODMR (Fig. S3). For that reason, we always used fresh samples for our experiments.

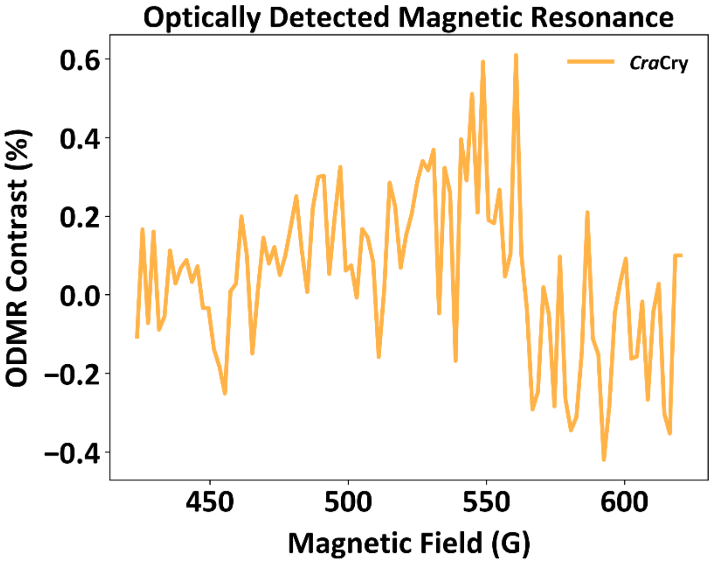


**Figure S3. Loss of the ODMR signal in *Cra*Cry samples after incubation at 37 °C for 16 hours.**

We also investigated *Drosophila melanogaster* cryptochrome (*Dm*Cry) samples, which exhibited strong and unexplained fluctuations in ODMR contrast. For that reason, we chose *Cra*Cry, where we could obtain highly reliable and stable ODMR signals.

**Supplementary Note 3. MFE laser power dependency iLOV**

Similar to the laser intensity dependence study performed on *Cra*Cry, iLOV was characterized. The strong enhancement of the MFE contrast with increasing laser power indicates that the excitation rate is a key factor in iLOV photochemical kinetics (Figure S4). This result indicates some mechanistic differences in the photochemical process of both proteins, which will be part of future studies.

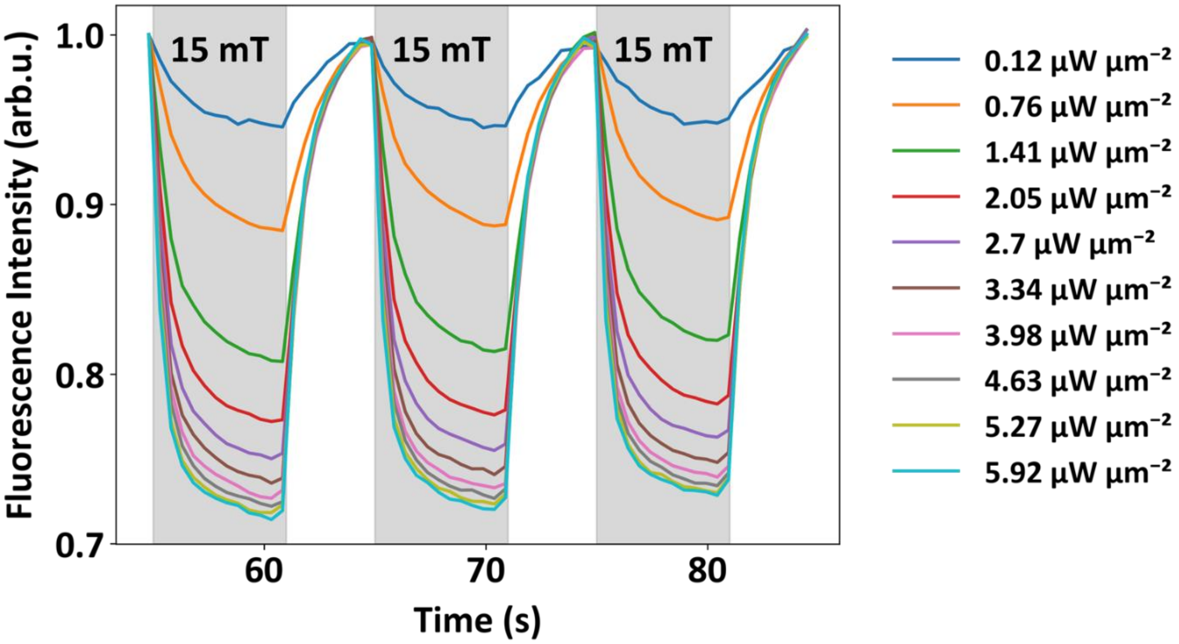


**Figure S4. MFE Laser power dependency iLOV.**

**Supplementary Note 4. Lineshape modulation induced by the RF delivery**

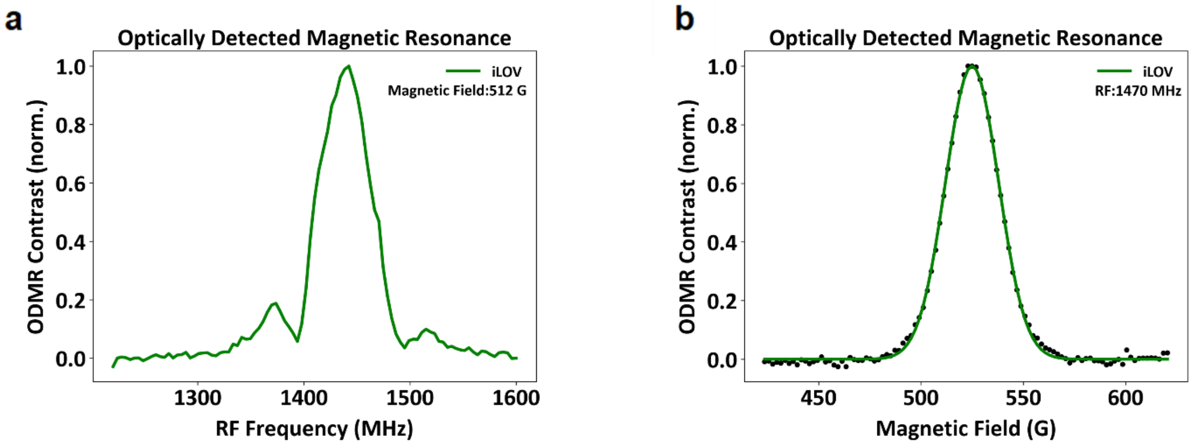


**Figure S5. Artifacts induced by the power dependency of the RF delivery.** a) An RF sweep at a static magnetic field shows a spectroscopic signature, which disappears when the magnetic field is swept under constant RF drive (b).

In Figure S5a, when the RF frequency is swept at a fixed magnetic field, the signal exhibits multiple peaks, which we attribute to uneven power delivery across the frequency band. To avoid this artifact, the RF frequency is fixed, and the magnetic field is swept (Fig. S5b). This configuration reveals a single peak, which is well-described by a Gaussian fit.

**Supplementary Note 5. Magnetic field strength calibration with boron nitride nanotubes (BNNTs)**

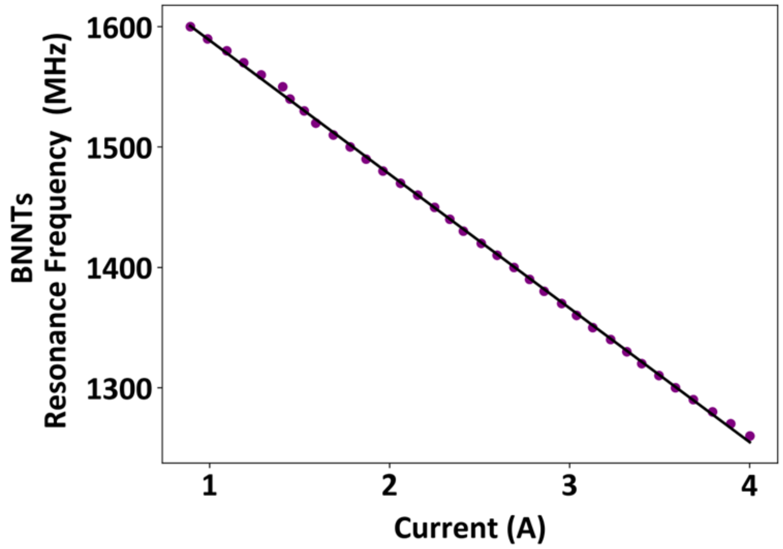


**Figure S6. Calibration of the current to magnetic field using ODMR of BNNTs**

The calibration of the magnetic field strength was performed using boron nitride nanotubes (BNNTs) purchased from NanoIntegris Inc. and have been characterized before[5,6]. The BNNTs have an average diameter of ~50 nm and lengths of a few micrometers. As-received BNNT powder was dispersed in 0.25% Triton X-100, stirred on a plate for 4 hours, followed by 10 minutes of sonication. Afterwards, the BNNT solution was applied to a glass coverslip in a laminar flow hood, allowing solution to evaporate and fix BNNTs on the coverslip. The laser power was adjusted to 500 µW to excite the BNNTs and fluorescence was collected using a 647 nm long pass filter (647 LP Edge Basic Longpass Filter, Semrock).

The calibration was performed by sweeping the current for the copper coils from 1 to 4 A while fixing the RF frequency at 35 distinct values, spaced evenly between 1250-1600 MHz. At each frequency, we recorded an ODMR spectrum of the BNNTs. Fitting each spectrum with a Gaussian function yielded the current for the resonance frequency. This procedure established a direct calibration curve between applied current and magnetic field strength.

**Supplementary Note 6. Optimizing ODMR contrast of iLOV**

To optimize the ODMR contrast, the RF frequency was set to 1256 MHz, the point of maximum contrast at 450 G. The fluorescence was recorded over 100 steps (10 ms exposure each) and the laser intensity was fixed at ~5.9 µW $\mu m^{-2}$. Figure S7a depicts fluorescence counts versus number of frames for “RF on” and “RF off”. Here, the readout time t constitutes a time trace of the fluorescence intensity evolution, where each step represents a 10 ms time point. In addition to the bleaching kinetics, the RF field produces a significant enhancement of fluorescence intensity. Analysis of the “RF on” / “RF off” fluorescence ratio (offset corrected) in Figure S7b reveals a steady increase in ODMR contrast as a function of readout time. These results indicate that the optimal ODMR contrast is obtained with the following sequence: simultaneous laser and RF excitation, followed by a delayed fluorescence readout which terminates synchronously with the laser turning off.

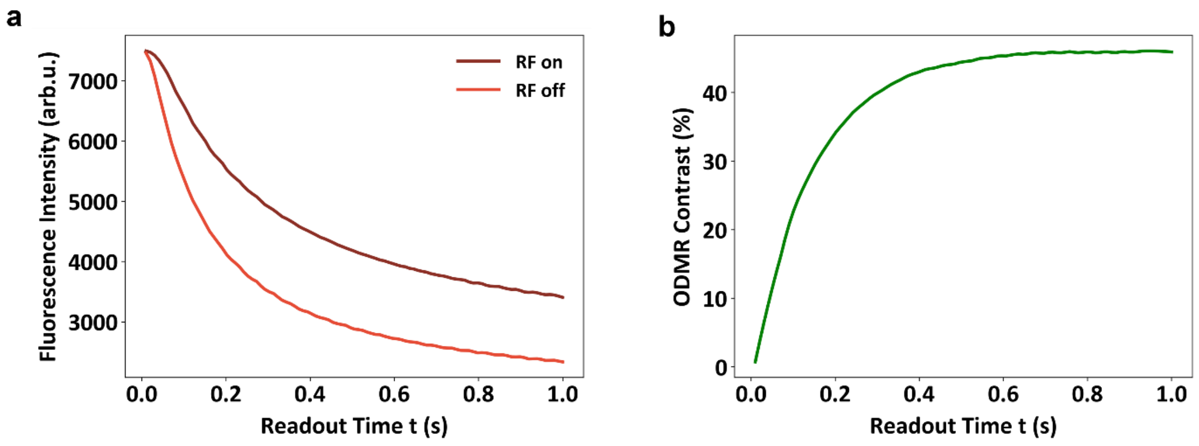


**Figure S7. Fluorescence intensity as a function of the readout time t (each step corresponds to 10 ms). a) Raw data. b) ODMR contrast.**

The second factor for optimizing ODMR contrast is laser intensity (see also Fig. S8). For this measurement, laser intensity was adjusted from ~ 0.1 to 5.9 μW $\mu m^{-2}$ (30 linear steps) and the exposure time was 500 ms. Each time a total of two frames were acquired, where the signal from the second frame was used for the analysis due to the larger contrast. Figure S8a demonstrates that the RF-induced fluorescence change increases with laser intensity. This is further quantified in Figure S8b by plotting the ODMR contrast ("RF on" / "RF off" -1). We attribute this to the fact that a higher laser intensity accelerates the photochemistry cycle of iLOV, allowing the system to reach the photochemical equilibrium faster.

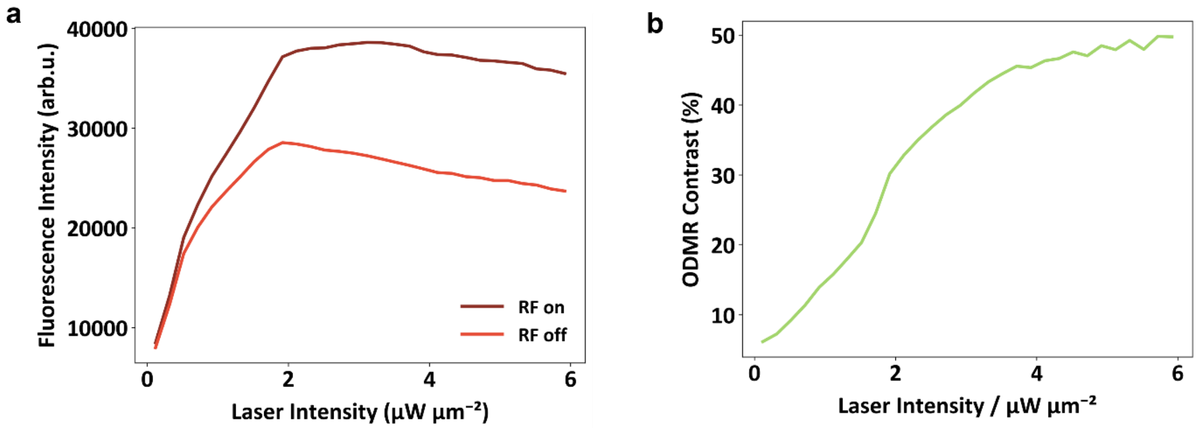


**Figure S8. Fluorescence intensity as a function of laser intensity. a) Raw data. b) ODMR contrast.**

These experiments demonstrate how RF and optical excitation (among other factors) affect the underlying chemical equilibrium of spin chemistry. This must be taken into account, for example, when performing more advanced pulsed ODMR experiments. In such experiments, the RF and optical excitation are separated (see also Supplementary Note 9), and any change in the excitation scheme will correspondingly lead to a different chemical equilibrium. This must be carefully considered when designing future experiments.

### Supplementary Note 7. ODMR spectra of iLOV recorded under different RF excitation frequencies

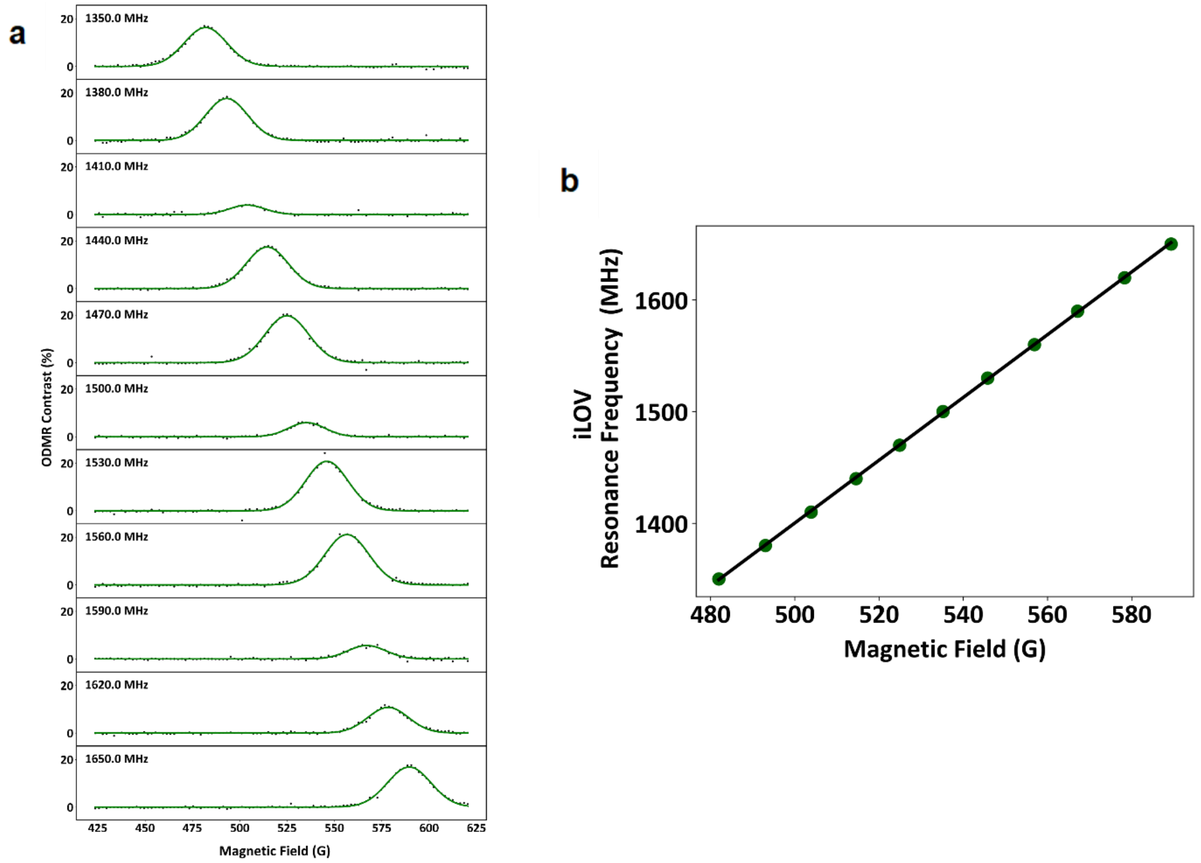


**Fig. S9. ODMR magnetic field resonance dependency on the RF excitation frequency**. a) ODMR spectra recorded under excitation at different static RF frequencies. b) ODMR resonance frequency as a function of the magnetic field strength, deduced from a fit of the data in a).

The iLOV ODMR spectrum was measured at selected RF frequencies. Sweeping the magnetic field strength shifts the ODMR resonance according to $g \approx 2$ ($\gamma_e$ = 2.81 ± 0.008 MHz/G). For the *Cra*Cry sample in Figure 2f we measure a gyromagnetic ratio of ($\gamma_e$ = 2.74 ± 0.05 MHz/G). The value deviates slightly from the ideal ones, which can be explained by the lower data quality of the *Cra*Cry dataset. Fluctuations of ODMR contrast observed across the frequency range are caused by the RF frequency dependent power delivery (mentioned in Supplementary Note 4).

### Supplementary Note 8. Stability and robustness of iLOV

To assess thermal stability, iLOV was kept at 37 °C for 16 hours. Subsequent ODMR measurements revealed no signal degradation, confirming iLOV's superior stability. We also performed experiments with the same sample over days at room temperature with only very minor degradation of the ODMR properties.

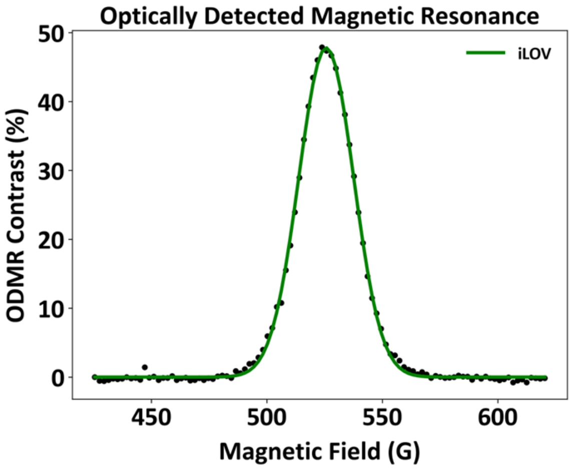


**Figure S10. iLOV ODMR after heating the sample in 37 °C for 16 hours.**

### Supplementary Note 9. Transient absorption spectroscopy and pulsed ODMR experiments

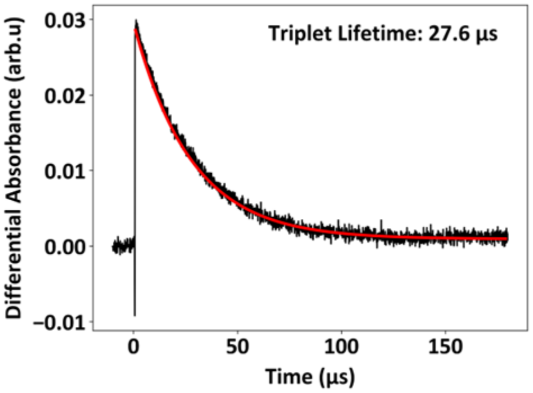


**Figure S11. Transient absorption spectroscopy of iLOV recorded at 600 nm.**

It is well known that LOV domains (incl. iLOV) form a triplet state of FMN after photoexcitation[7]. For that reason, we performed transient absorption (TA) experiments to probe the triplet state lifetime for our sample. Figure S11 shows the room temperature TA decay of iLOV, probed at 600 nm after 460 nm excitation. A single-exponential fit yields a triplet-state FMN lifetime of ~27.6 µs. This lifetime is in agreement with results obtained from Kopka et al.[7] and we assign this lifetime to the decay of the $^3$FMN state accordingly. TA did not provide any direct spectroscopic evidence for a radical pair, suggesting that it might be obscured by the dominant triplet signal or only weakly populated.

These experiments were performed on a commercially available laser flash photolysis spectrometer (LP920K, Edinburgh Instruments Ltd., Livingston, GB). Sample excitation was achieved using a Nd:YAG laser (Surelite II, Continuum, Santa Clara, CA, USA) with an internal pump frequency of 10 Hz. Using optical generators for the second and third harmonic, emission was set to 355 nm at a duration of 6 ns and a pulse energy of approx. 100 mJ. These pulses pumped an optical parametric oscillator (Continuum OPO PLUS, Continuum, Santa Clara, CA, USA). Samples were excited at 460 nm with a pulse energy of (6.0 ± 1.0) mJ and probed inside a quartz cuvette (H108F-QS, Hellma Analytics, Müllheim, Germany) at a temperature of (292.0 ± 0.1) K using a flow through thermostat (LAUDA Alpha RF8, Lauda Dr.

R. Wobser GmbH & Co. KG, Lauda-Königshofen, Germany). Irradiated samples were probed with a xenon shortarc lamp (XBO450W, Osram, Graz, Austria) with a pulse length of 10 ms. Detection was performed using a multi-channel plate and digitalized using an oscilloscope (TDS-3012, Tektronix Inc., Beaverton, Oregon, USA) with a maximum time resolution of 4 ns. Final measuring frequency was set to ~0.2 Hz, equaling a delay of 5 s between measurements.

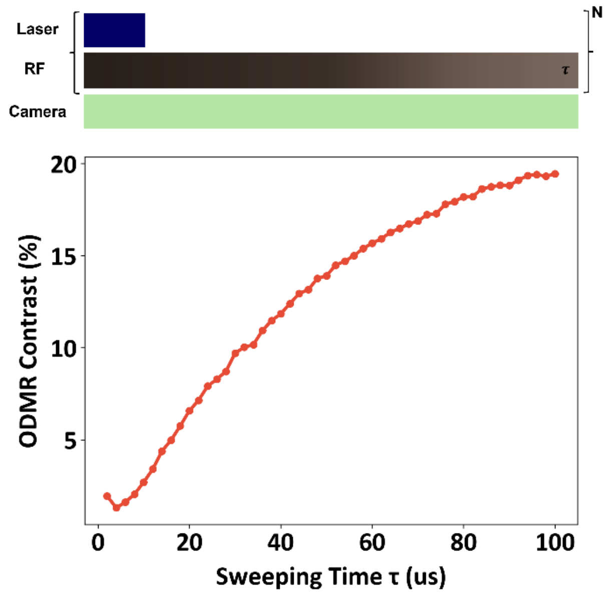


**Figure S12. Pulsed ODMR experiments.**

The ODMR experiments presented in the main manuscript were performed under continuous (cw) RF and optical excitation (apart from the on/off cycles required for sample recovery). To gain mechanistic insight, we separated the optical excitation from the RF control (pulsed ODMR). This approach provides, for instance, information on how long the spin state can be manipulated after optical excitation. In Figure S12, the excitation laser is pulsed with a fixed 10 μs duration (10% duty cycle compared to the cw ODMR experiments), while the RF pulse duration is swept from 2 μs to 100 μs. The laser-RF pulse cycle was iterated continuously over a total camera acquisition period of 100 ms. The reduced duty cycle (i.e. shorter laser exposure and consequently fewer photons per experiment) leads to a reduction in the overall ODMR contrast (see also Supplementary Note 6). The ODMR contrast continues to increase with increasing RF pulse duration, persisting for tens of microseconds after optical excitation—comparable to the lifetime of the FMN triplet state in iLOV (we note that the pulse ODMR and TA experiments have been performed at different magnetic fields, which may slightly impact the lifetimes).

The SCRP lifetime and the spin polarization of the FMN triplet are both very likely much shorter than this time scale. In a simple model where the FMN triplet state decays both to the FMN ground state (e.g., via phosphorescence) and to a SCRP, the overall triplet decay rate is the sum of the two individual rates ($k_{GS} + k_{SCRP}$). This is the same rate at which the SCRP is formed, namely $k_{GS} + k_{SCRP}$. Consequently, the FMN triplet state (with a lifetime of tens of microseconds) continuously generates SCRPs throughout its lifetime, and this is exactly what is observed in Figure S12.

### Supplementary Note 10. Linewidth analysis for iLOV

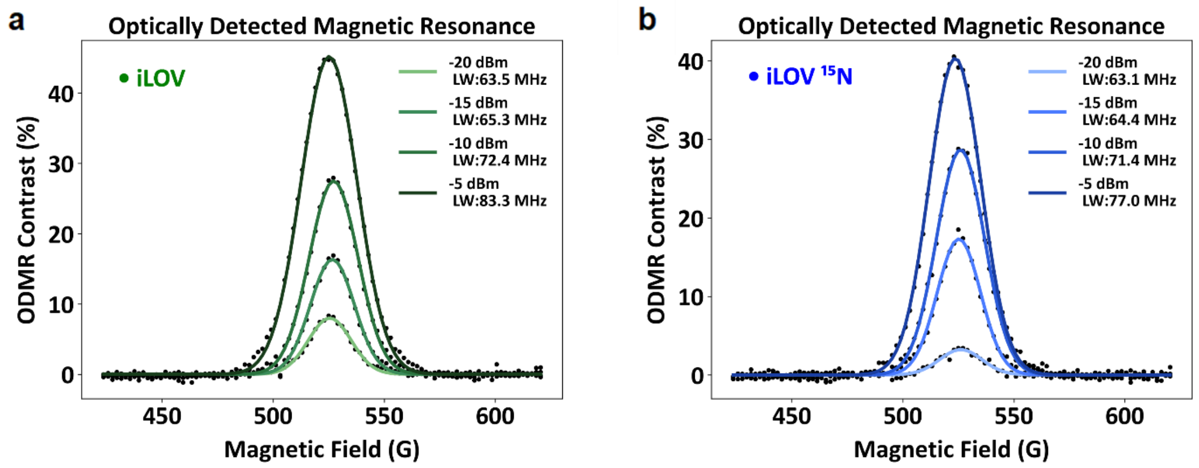


**Figure S13. ODMR signal as a function of RF power.** The power values refer to the output power of the signal source and do not reflect the actual power at the sample position. a) iLOV ODMR signal as a function of RF power b) Uniformly labeled $^{15}N$ iLOV ODMR signal as a function of RF power.

To avoid artifacts from RF-power broadening, we performed linewidth measurements as a function of the applied RF power (Fig. S13). We cannot determine the exact RF power at the sample position; therefore, we report the output power of our signal generator (which is subsequently amplified). We observe that the linewidth reaches a minimum below –15 dBm, indicating that in this regime the linewidth is not power-dependent. Both samples, the natural-abundance material and the $^{15}N$-labelled version, show highly similar behavior, demonstrating that the $^{15}N$ hyperfine interaction does not introduce a significant influence.
For the *Cra*Cry sample, higher RF power was required; otherwise, the ODMR signal would have been too weak. For that reason, we used an output power of -10 dBm at the signal source for the comparison measurements shown in Figure 3.

As described in the Methods section, CraCry and iLOV were initially prepared in different buffer solutions. To rule out a potential solvent effect, notably the glycerol concentration, on the ODMR linewidth, we measured iLOV in the *Cra*Cry buffer (50 mM $NaH_2PO_4$ pH 7.8, 100 mM NaCl, 50% (v/v) glycerol). The result shows no significant change in linewidth, confirming that the observed spectral differences are intrinsic to the proteins and not an artifact of the solvent environment.

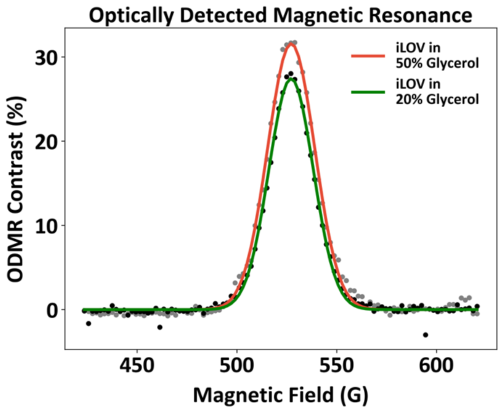


**Figure S14. ODMR signal of iLOV in two different solvent environments.**

**Supplementary Note 11. Magnetic field calibration with RF frequency sweeping**

For the magnetic field imaging shown in Figure 4, we chose to sweep the RF resonance frequency under static magnetic field conditions. This comes with the artifacts induced by the RF frequency dependent lineshape as discussed in Supplementarty Note 4. Figure S15a depicts the iLOV ODMR spectrum at two different magnetic fields: the light green curves correspond to the lower field (440 G), and the dark green curves to the higher field (449 G) with a strong modulation in the lineshape. We found that we can empirically estimate the average resonance frequency using

$$\overline{f} = \frac{\sum_i freq_i \cdot C_i}{\sum_i C_i}$$

Where $\overline{f}$ is the average resonance frequency, $freq_i$ is the swept RF frequency and $C_i$ is the ODMR contrast for each RF frequency. The estimated average resonance frequencies are marked by a red star (for the light green data) and a blue star (for the dark green data) in Figure 15b which corresponds very well with the BNNTs resonances (black and grey) at these magnetic fields.

We further investigate the accuracy of average resonance frequency estimation in Figure S15c. Using five ODMR spectra from iLOV and BNNTs, the average resonance frequency of iLOV was fitted as a linear function of the calibrated magnetic field strength. The resulting slope closely matches the electron gyromagnetic ratio of ~ 2.8 MHz/G.

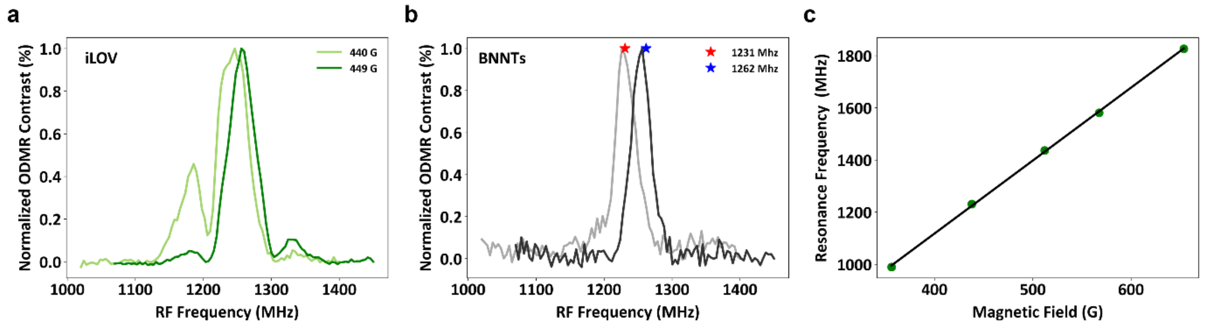


**Figure S15. Magnetic field calibration of iLOV for imaging applications.** a) and b) ODMR spectra at different magnetic fields of iLOV and BNNTs, respectively. c) Calibration with our empirical model.

**Supplementary Note 12. Validation of the magnetic field gradient with BNNTs**

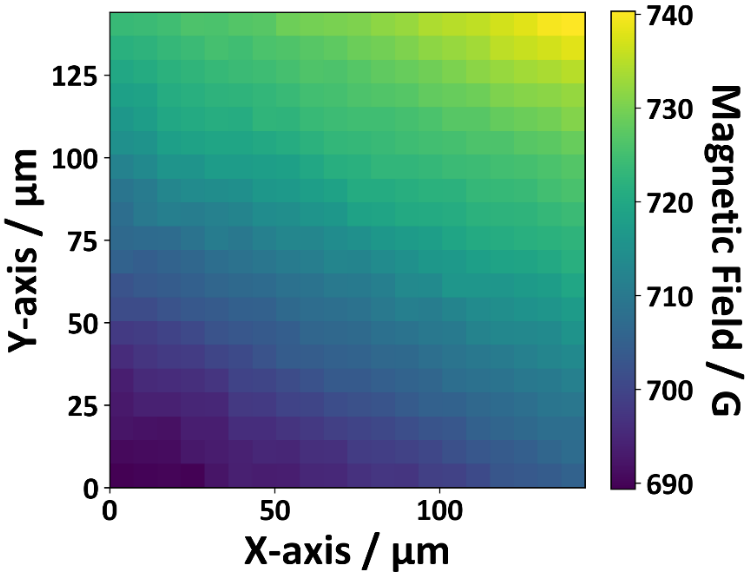


**Figure S16. Magnetic field imaging using BNNTs.**

To validate the magnetic field map shown in Figure 4b, we used BNNTs homogeneously dispersed on a glass plate as a reference, positioned in the same way as the protein sample. Magnetic field imaging of the BNNTs reveals the overall same spatial gradient (decreasing from the upper right to the lower left) that we observe for the iLOV system. The slightly steeper gradient in the BNNT data may result from the manual positioning of the samples relative to the magnet and the associated differences in their exact placement. Additional deviations may arise from differences in sample geometry: the BNNT sample forms a thin solid film (< 10 µm), whereas the iLOV sample is a ~100 µm-thick liquid layer trapped between cover slides.

We also note that, due to the excitation-dependent ODMR contrast (Supplementary Note 6) of the protein systems, the Gaussian excitation profile of the laser leads to reduced SNR at the corners of the image. This issue can be mitigated in future experiments by shaping the laser beam profile.
